# Genetic analysis and mapping of adult plant stripe rust resistance loci in CIMMYT wheat ‘Kijil’ under Mexican and Chinese field environments

**DOI:** 10.64898/2026.01.28.702223

**Authors:** Shanshan Yan, Lichao Teng, Menghan Xi, Chan Yuan, Liang Wang, Shunda Li, Julio Huerta-Espino, Sridhar Bhavani, Ravi P. Singh, Caixia Lan

## Abstract

Stripe rust, caused by *Puccinia striiformis* f. sp. *tritici*, can cause severe yield losses in wheat (*Triticum aestivum* L.) during epidemics. Breeding resistant wheat varieties remains the most cost-effective approach to manage this disease; and the identification of new resistance loci is essential for maintaining genetic diversity. The CIMMYT-derived wheat line ‘Kijil’ was highly resistant to stripe rust in both Mexican and Chinese environments. A population of 153 F₅ recombinant inbred lines (RILs) was derived from a cross between Kijil and the susceptible parent ‘Apav#1’. The population was phenotyped for stripe rust resistance across seven environments in two countries and genotyped using a genotyping-by-sequencing (GBS) platform. Inclusive composite interval mapping (ICIM) was uesd to construct a genetic map and identify significant resistance quantitative trait loci (QTLs) using 5,468 polymorphic markers. Mapping revealed the known resistance loci *Yr29*, *Yr30* and *QYr.hzau-3AS*, along with two novel loci, *QYr.hzau-2BS* and *QYr.hzau-5DL*, across both Chinese and Mexican rust environments. Among these, *QYr.hzau-2BS* accounted for 11.75% to 19.19% of the phenotypic variance. A corresponding KASP marker, KASP_2BS, was developed to facilitate maker-assisted selection. Based on the mapping interval, four candidate genes underlying this locus were predicted. Further analysis revealed that *Yr29* showed significant additive effects with other stripe rust resistance genes/loci, and the combination of *Yr29*, *Yr30*, and *QYr.hzau-2BS* reduced disease severity by up to 67.8%. Our findings suggest that Kijil and RILs carrying *Yr29*, *Yr30,* and *QYr.hzau-2BS* can serve as valuable donors for breeding wheat varieties with improved stripe rust resistance.

**Author summary:** Stripe rust is an important disease that seriously threatens the yield and quality of wheat. It is crucial to explore new resistant resources and cultivate durable resistant varieties in the current breeding programme. In this study, we analysed the genetic basis of resistance to stripe rust in the wheat line “Kijil”, which has broad-spectrum resistance to stripe rust. Through genetic mapping, we identified five quantitative trait loci for stripe rust, including two new resistance loci. A closely linked KASP marker, KASP_2BS, was developed for the *QYr.hzau-2BS*, which can be used for rapid and accurate screening of resistant plants in early breeding populations. Meanwhile, within this QTL region, we screened four candidate genes based on expression analysis. In addition, it was found that polymerization of *QYr.hzau-2BS* with known resistance genes, *Yr29* and *Yr30*, significantly enhanced resistance and reduced disease severity to low levels and near immunity. In conclusion, this study provides new genetic resources, practical molecular markers and effective gene polymerization strategies for breeding wheat for stripe rust resistance. ‘Kijil’ and the lines containing resistance loci have important breeding utilization value.

## Introduction

Wheat is one of the world’s most important food crops, providing a major source of energy and protein for humans [1] and accounting for about 30% of global cereal consumption (FAO, https://www.fao.org). However, wheat yields and quality are seriously threatened by rust epidemics [2-4]. Stripe, or yellow rust (YR), caused by *Puccinia striiformis* f. sp. *tritici* (*Pst*), is one of the most destructive wheat diseases worldwide. It is estimated that wheat rust causes global economic losses amounting to billions of US dollars annually [5]. The frequency and severity of rust outbreaks have increased in recent years, driven by the emergence of highly virulent races and changing climatic conditions [6]. Developing and deploying new, durable sources of resistance is therefore one of the most economical and environmentally sustainable strategies for disease management.

Resistance to YR in wheat can be broadly categorized into two types: seedling resistance or all-stage resistance (ASR), and adult plant resistance (APR). ASR, also known as vertical resistance, is typically conferred by single, major-effect genes that are expressed throughout the plant’s life cycle [7]. However, this type of resistance is often short-lived in the field due to pathogen evolution and the emergence of new virulent races [8]. Large-scale deployment of cultivars with ASR genes has repeatedly led to yield losses due to the presence of new virulent *Pst* races resulting in the breakdown of resistance genes, such as *Yr9*, *Yr10*, and *Yr24/Yr26* in China [9, 10]. In contrast, APR—also referred to horizontal resistance—is controlled by multiple minor-effect genes and is usually expressed at later growth stages [11]. APR is generally considered more durable, as it tends to be non-race specific. Several well-known APR genes, such as *Lr34/Yr18* [12], *Lr46/Yr29* [13], *Lr67/Yr46* [14], and *Sr2/Yr30* [15], have provided effective resistance to multiple diseases for over a century. To date, 87 stripe rust resistance genes have been officially catalogued, and more than 406 QTLs associated with stripe rust resistance have been mapped to the wheat reference genome [16, 17], including contributions from over 20 wheat-related species. Given the limited availability of resistance genes in cultivated wheat and the ongoing evolution of new virulence, the discovery and strategic combination of new resistance loci are still the key for stripe rust management.

The CIMMYT-derived wheat line ‘Kijil’ was susceptible at the seedling stage to both Mexican and Chinese *Pst* races but showed high levels of APR under field conditions. In this study, we developed 153 F₅ recombinant inbred lines (RILs) from a cross between the susceptible cultivar ‘Apav#1’ and Kijil. The objectives of this study were to: (1) elucidate the genetic basis of APR to YR in Kijil; (2) identify new resistance loci using molecular markers under multi-environment tests and develop molecular markers associated with the newly identified loci for wheat breeding programs; and (3) assess the interaction among detected resistance loci under both Chinese and Mexican field environments.

## 2. Results

### 2.1. Phenotypic analysis

At the seedling stage, the infection types (IT) of Apav#1 and Kijil were 7-8 and 6-7, respectively, against *Pst* races CYR33 and CYR34. At the adult plant stage, Apav#1 showed DS ranging from 40% to 100% with susceptible (S) or moderately susceptible (MS) reactions across seven test environments, whereas Kijil displayed 1MS and 1MR reactions at Toluca and Batan, Mexico, respectively (Fig 1A). In Chinese YR nurseries, Kijil exhibited FDS and host reactions of 0–1R, the mean DS across the RIL population ranged from 22.04% to 51.93% (Fig 1A and S1 Fig); in Mexican environments, the mean DS across the RIL population ranged from 16.35% to 38.86% ((Fig 1A and S1 Table). The unbiased linear predicted values of phenotypic data from the Mexican (BLUPM) and Chinese environments (BLUPC) had a mean value of 37.46% and 37.18%, respectively, and both approximately followed a normal distribution (Fig 1B). In all test environments, the frequency distribution of RILs for YR severity was continuous, indicating that APR was quantitatively inherited in this population. Mendelian segregation analysis of observed and expected frequencies in three phenotypic classes suggested the presence of two to five resistance genes with additive effects, depending on the test environment (Table 1).

**Fig 1.**
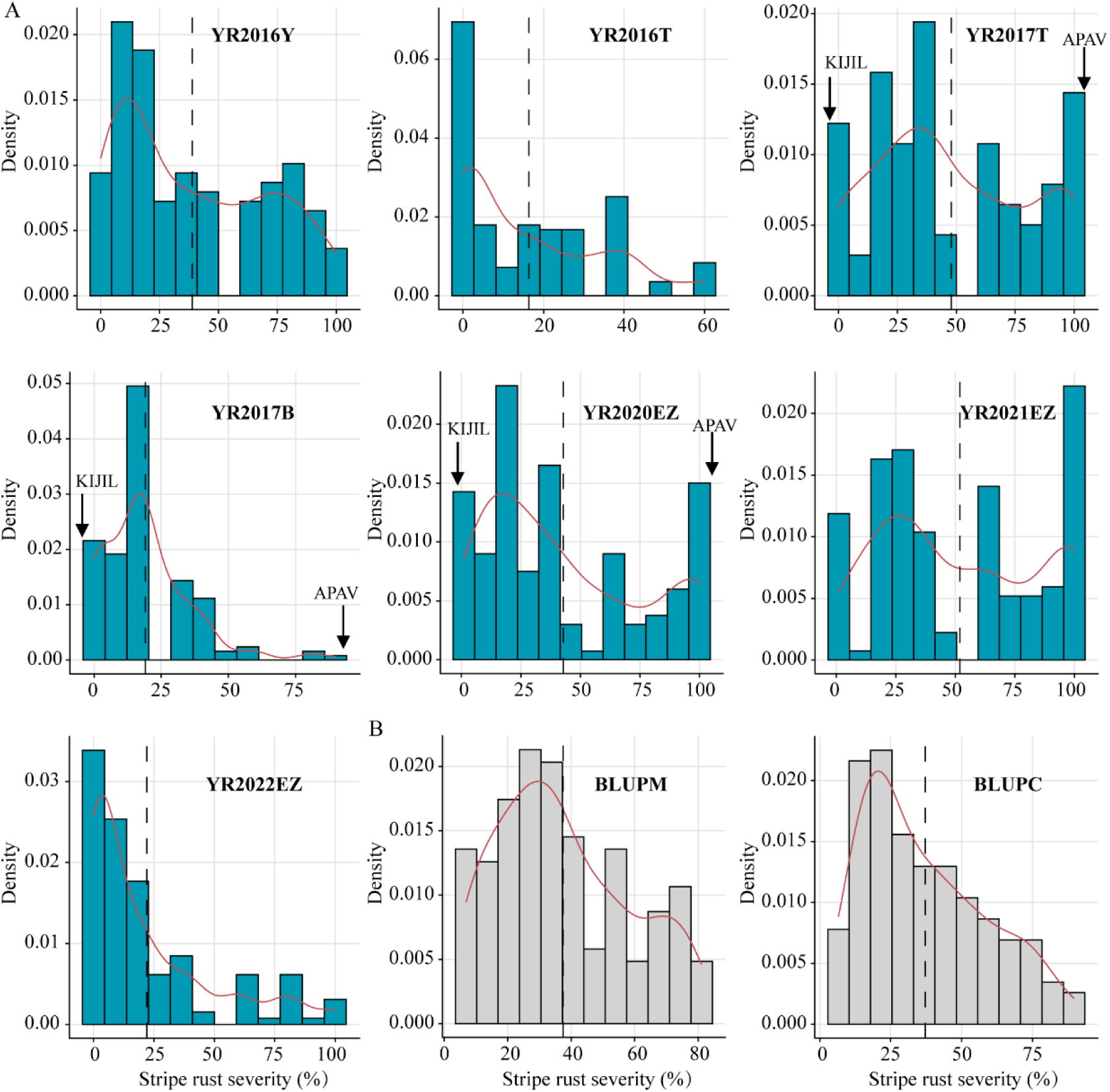
Frequency distribution of stripe rust severity in a wheat RIL population under different field environments. Frequency distributions for final disease severity (FDS) of stripe rust for 153 ‘Apav#1’×‘Kijil’ F_5_ recombinant inbred lines (RILs) in field trials at Obregón, Mexico, during 2015-2016 (YR2016Y) and at Toluca, Mexico, during 2015-2016 (YR2016T) and 2016-2017 (YR2017T) and at Batan, Mexico, during 2016-2017 (YR2017B); FDS for stripe rust at Ezhou, Hubei Province, China, during 2019-2020 (YR2020EZ), 2020-2021 (YR2021EZ) and 2021-2022(YR2022EZ). **(B)** Frequency distributions of the best linear unbiased prediction (BLUP) values for stripe rust severity. BLUPM and BLUPC: the unbiased linear predicted values of phenotypic data from the Mexican and Chinese environments, respectively. The vertical dashed black lines represent the mean disease severity for each environment.

**Table 1.** Estimated number of resistance genes that conferred adult plant resistance to stripe rust based on the Mendelian segregation ratios in ‘Apav#1’ × ‘Kijil’ recombinant inbred lines population.

| Category | YR2016Y | YR2016T | YR2017T | YR2017B | YR2020EZ | YR2021EZ | YR2022EZ |
| --- | --- | --- | --- | --- | --- | --- | --- |
| HTPR | 26 | 13 | 6 | 27 | 21 | 16 | 2 |
| HPTs | 10 | 5 | 20 | 22 | 29 | 30 | 4 |
| Others | 117 | 134 | 127 | 104 | 98 | 104 | 137 |
| Total | 153 | 152 | 153 | 153 | 148 | 150 | 143 |
| No. of genes | 3 | 4 | 3 | 2 | 2 | 2 | 5 |
| P-value<sup>a</sup> | 0.00* | 0.81 | 0.02* | 0.23 | 0.30 | 0.03* | 0.50 |

HTPR, homozygous parental type resistant (RILs showing a similar phenotype as the resistant parent); HPTs, homozygous parental type susceptible (RILs showing a similar phenotype as the susceptible parent); Others, RILs showing different responses from the above two categories; YR2016Y, final stripe rust severity, Ciudad Obregon (Mexico) 2015-2016; YR2016T, final stripe rust severity, Toluca (Mexico) 2016; YR2017T final stripe rust severity, Toluca (Mexico) 2017; YR2017B, final stripe rust severity, Batan, (Mexico) 2017; YR2020EZ, final stripe rust severity, Ezhou (China) 2020; YR2021EZ, final stripe rust severity, Ezhou (China) 2021; YR2022EZ, final stripe rust severity, Ezhou (China) 2022.
<sup>a</sup> Asignificant *p* value (\*) indicates deviation from the expected ratio at *P* < 0.05.

Pearson correlation coefficients of FDS among RILs ranged from 0.67 to 0.83 in Mexican environments and 0.57 to 0.71 in Chinese environments (Table 2). In addition, FDS was significantly correlated across all test environments, with correlation coefficients ranging from 0.49 to 0.66. These relatively high correlations between Mexican and Chinese trials indicated that Kijil conferred broad resistance to *Pst* across both the environments.

**Table 2.** Phenotypic Pearson’s correlations for stripe rust in the 153 ‘Apav#1’× ‘Kijil’ F_5_ RIL population using the final disease scores (FDS) in each environment.

| Environment | YR2016T | YR2016Y | YR2017T | YR2017B | YR2020EZ | YR2021EZ |
| --- | --- | --- | --- | --- | --- | --- |
| YR2016Y | 0.70** |  |  |  |  |  |
| YR2017T | 0.83** | 0.72** |  |  |  |  |
| YR2017B | 0.70** | 0.67** | 0.81** |  |  |  |
| YR2020EZ | 0.65** | 0.59** | 0.63** | 0.55** |  |  |
| YR2021EZ | 0.61** | 0.66** | 0.61** | 0.52** | 0.70** |  |
| YR2022EZ | 0.61** | 0.65** | 0.54** | 0.49** | 0.57** | 0.71** |

YR2016Y, final stripe rust severity, Ciudad Obregon (Mexico) 2015-2016; YR2016T, final stripe rust severity, Toluca (Mexico) 2016; YR2017T final stripe rust severity, Toluca (Mexico) 2017; YR2017B, final stripe rust severity, Batan, (Mexico) 2017; YR2020EZ, final stripe rust severity, Ezhou (China) 2020; YR2021EZ, final stripe rust severity, Ezhou (China) 2021; YR2022EZ, final stripe rust severity, Ezhou (China) 2022. \*\* Significant at $P < 0.01$ .

### 2.2. Genetic linkage maps

A total of 5,468 polymorphic markers, comprising 4,480 PAV markers, 984 SNP markers, 3 SSR markers and 1 KASP marker, were assigned to 42 linkage groups distributed across the A, B, and D genomes, with 1,826, 3,163, and 479 markers, respectively (S1 File). The genetic map spanned 8,523.94 cM, with an average marker density of 1.55 cM per marker. Linkage maps harboring identified QTLs are shown in the present study (S1 File).

### 2.3. Mapping of adult plant stripe rust resistance loci

Five QTLs for YR resistance, *QYr.hzau-1BL*, *QYr.hzau-2BS*, *QYr.hzau-3AS*, *QYr.hzau-3BS*, and *QYr.hzau-5DL*, were detected on chromosomes 1BL, 2BS, 3AS, 3BS, and 5DL, respectively, using ICIM software based on 1,000 permutations. All QTLs were contributed by the resistant parent Kijil (Table 3). Among these, *QYr.hzau-1BL*, located on chromosome 1BL, was consistently detected in 9 environments (YR2016Y, YR2016T, YR2017T, YR2017B, YR2020EZ, YR2021EZ, YR2022EZ, BLUPM and BLUPC), explaining 16.03–33.71% of the phenotypic variation (Table 3; Fig. 2A). Based on the physical positions of flanking markers, this QTL corresponds to the known resistance gene *Yr29*. *QYr.hzau-2BS* was detected in 7 environments (YR2016Y, YR2016T, YR2020EZ, YR2021EZ, YR2022EZ, BLUPM and BLUPC), explaining 5.03–23.39% of the variation. It was closely linked to markers 1228364 and 3021343 (Table 3; Fig. 2B) and showed no linkage to the known stripe rust resistance gene *Yr27*. *QYr.hzau-3AS*, flanked by markers 1176628 and 981733, explained 5.19–17.53% of the variation and was only detected in Mexican environments (Table 3; Fig. 2C). *QYr.hzau-3BS*, located on chromosome 3BS between markers 1321522 and *Xgwm533*, explained 3.48–12.77% of the variation in three environments (YR2016T, YR2017B, YR2020EZ, YR2021EZ, YR2022EZ, BLUPM and BLUPC) (Table 3; Fig. 2D), and likely corresponds to *Yr30*. *QYr.hzau-5DL* was detected in three Mexican environments and two Chinese environment between markers 1102087 and 3026417, explaining 4.15–17.40% of the phenotypic variation (Table 3; Fig. 2E).

**Fig 2.**
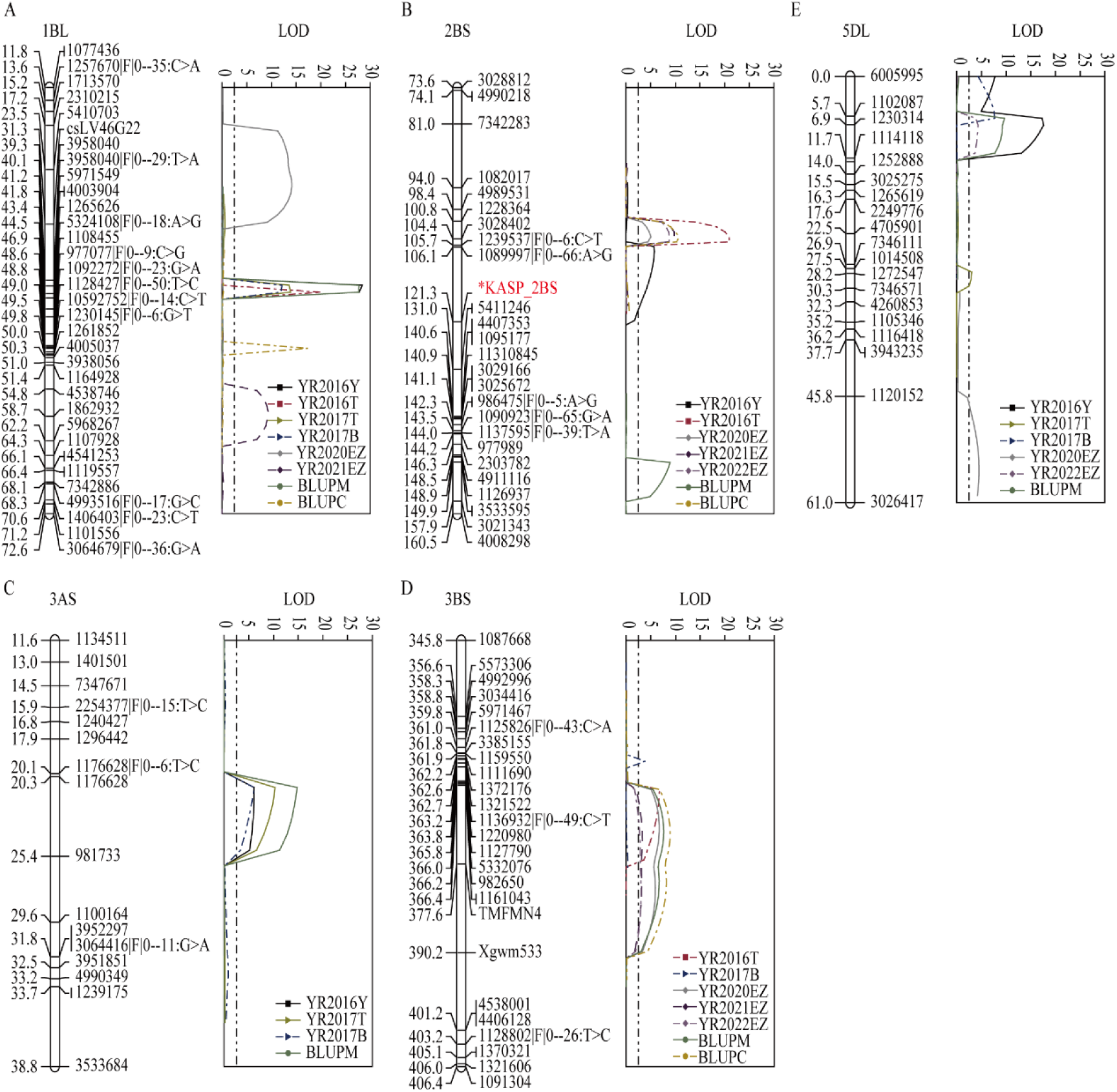
Quantitative trait loci (QTL) likelihood plots for stripe rust on five wheat chromosomes. **(A)** chromosome 1BL. **(B)** chromosome 2BS. **(C)** chromosome 3AS. **(D)** chromosome 3BS. **(E)** chromosome 5DL. The significant LOD threshold was based on 1,000 permutations. Positions in cM of the molecular markers along chromosomes are shown on the vertical axes using cumulated genetic distances. YR2016Y: phenotypic data for stripe rust resistance recorded at Ciudad Obregon, Mexico, during 2016-2017 crop season; YR2016T, YR2017T and YR2017B: phenotypic data for stripe rust resistance recorded at Toluca and Batan, Mexico, during 2016 and 2017 respectively; YR2020EZ, YR2021EZ, YR2022EZ: phenotypic data for stripe rust resistance recorded at Ezhou, China, during 2019-2020, 2020-2021 and 2021-2022 crop seasons; BLUPM and BLUPC : the unbiased linear predicted values of phenotypic data from the Mexican and Chinese environments, respectively.

**Table 3.**
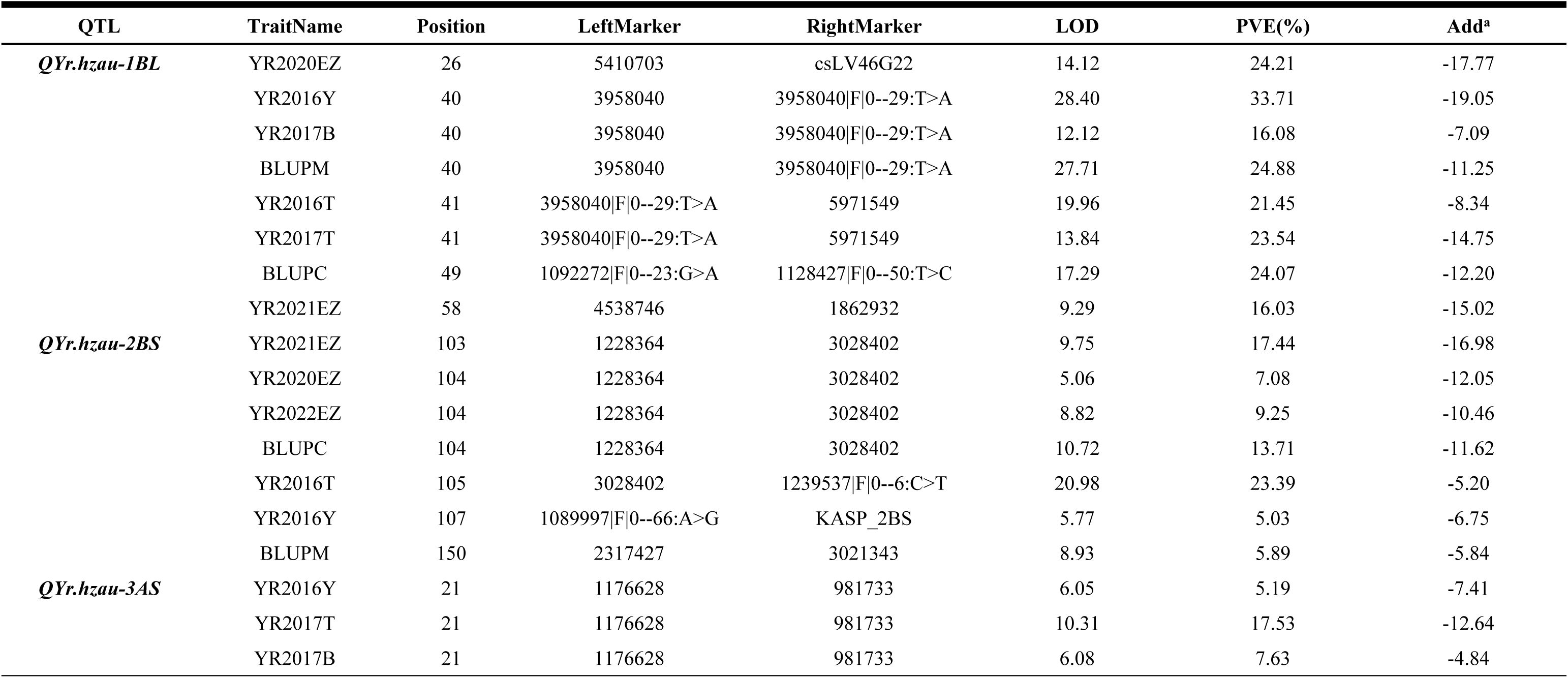
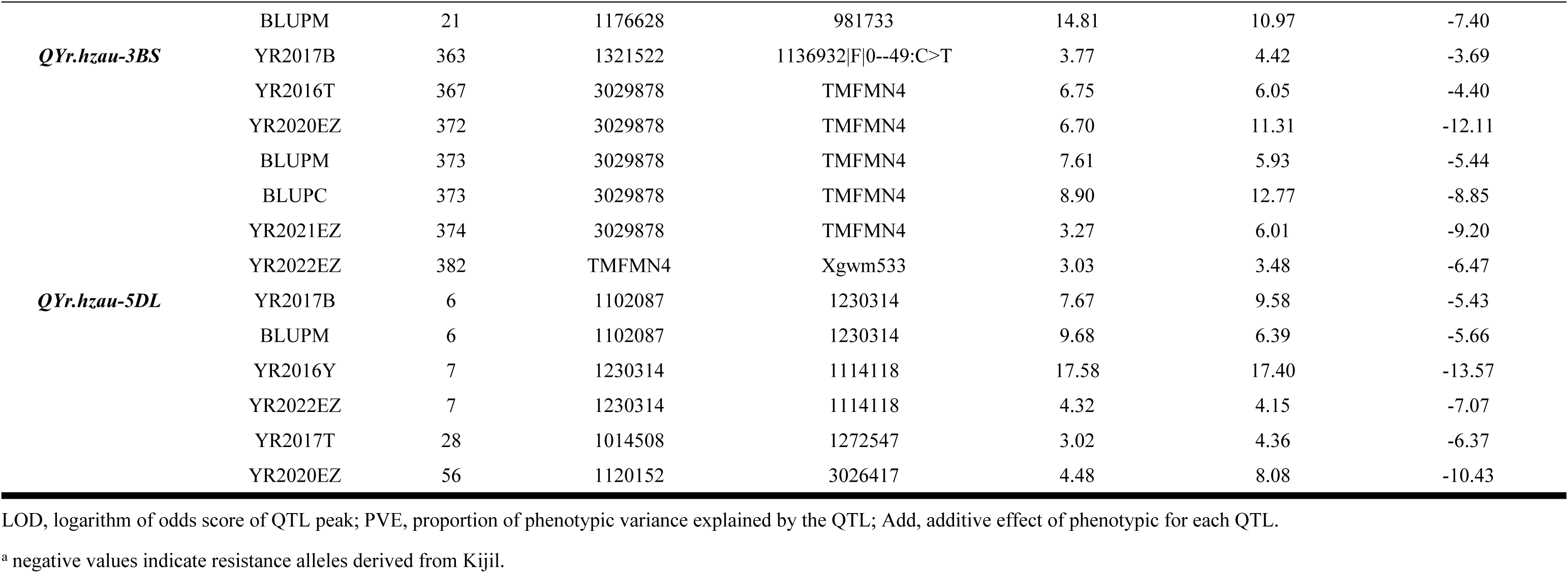
Position and effect of quantitative trait loci (QTL) detected for adult plant resistance (APR) to stripe rust (YR) in both Mexican and Chinese rust environment, using final disease severity and best linear unbiased prediction (BLUP; BLUPM for Mexico, BLUPC for China)) of 153 RILs derived from the cross of ‘Apav#1’× ‘Kijil’.

### 2.4. Development and validation of KASP molecular marker for *QYr.hzau-2BS*

Considering the stable phenotypic effect of *QYr.hzau-2BS* across multiple environments, we developed a Kompetitive Allele-Specific PCR (KASP) marker based on the closely linked SNP marker 1239537|F|0--6:C>T and it was named KASP_2BS (S2 Table). Genotyping results showed clustering of the alleles into two distinct groups (Fig 3A). Linkage analysis confirmed that KASP_2BS is located precisely at the peak of the QTL and it was placed to the physical position of 134.55 Mb on chromosome arm 2BS based on the Chinese Spring RefSeq v1.0 genome. We used this marker to genotype the entire RIL population; and it was significantly associated with disease severity reductions in one Mexican environment (YR2016T) and four Chinese environments (YR2020EZ, YR2021EZ, YR2022EZ, and BLUPC), with reductions of 8.89%, 12.61%, 18.13%, 13.64%, and 11.72%, respectively (Fig 3B). Furthermore, this marker also showed disease severity reductions of 3.00%, 5.13%, 4.92%, and 5.19%, respectively, under YR2016Y, YR2017T, YR2017B, and BLUPM rust environments (S2 Fig).

**Fig 3.**
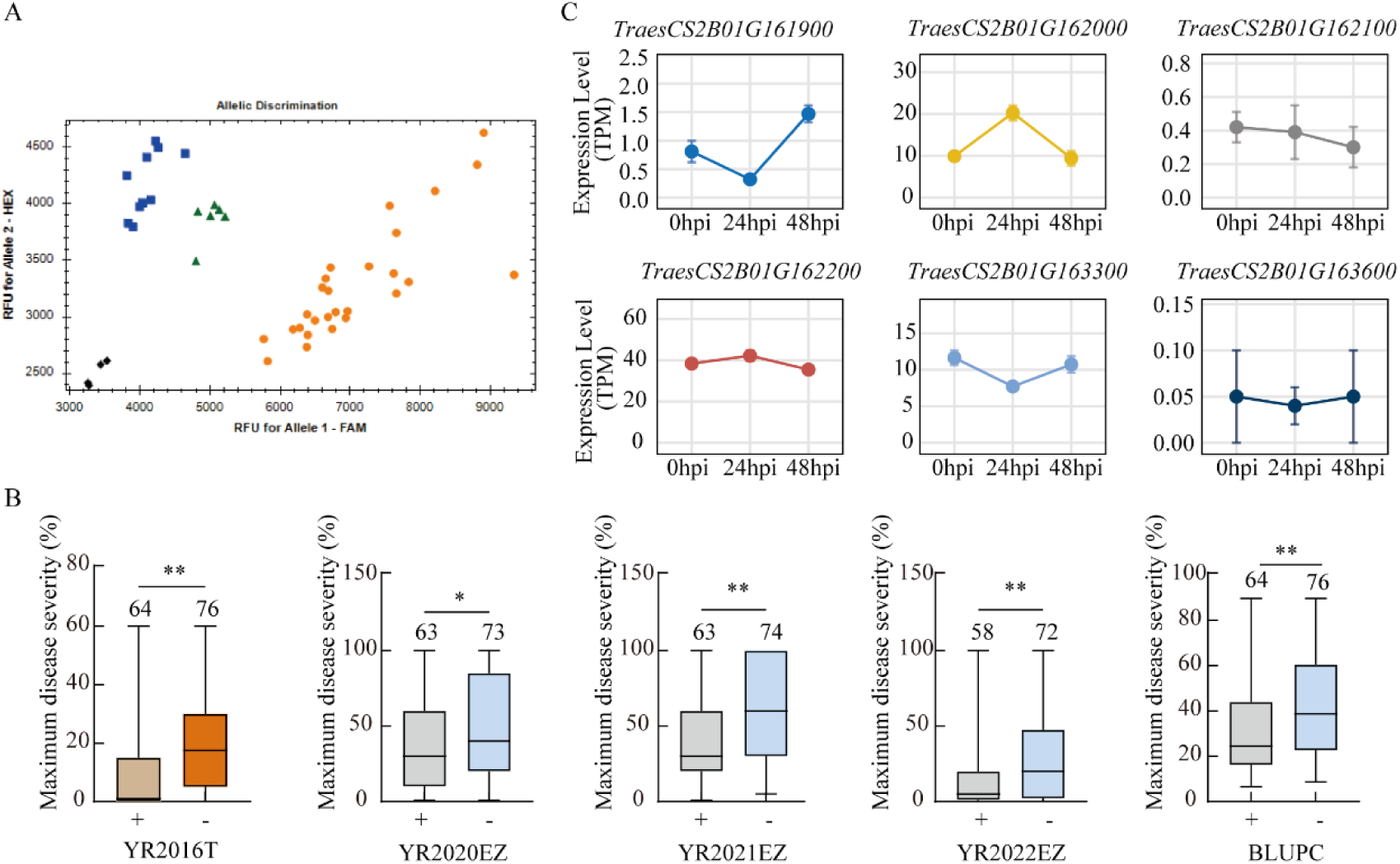
Development of a KASP Marker and functional characterization of candidate genes for *QYr.hzau-2BS*. (**A**) KASP genotyping for *QYr.hzau-2BS*. Blue, orange, and green dots represent susceptible, resistant and heterozygous genotypes, respectively; black dots indicate no template control. (**B**) Association analysis for *QYr.hzau-2BS* using the KASP marker with stripe rust disease severity across five environments (YR2016T, YR2020EZ, YR2021EZ, YR2022EZ and BLUPC). “−” indicates the absence of the QTL locus, whereas “+” indicates its presence. Significance levels are denoted by asterisks (*, *P* < 0.05; **, *P* < 0.01; ns = non-significant). The sample size was designed. (**C**) Expression profiles (transcripts per million, TPM) of six candidate genes within the *QYr.hzau-2BS* interval in the Avocet+*Yr5* line at 0-, 24-, and 48-hours post-inoculation (Hpi) with *Puccinia striiformis* f. sp. *tritici* (n=3). Error bars represent ± SEM.

Based on linkage marker analysis, 19 annotated genes were identified within the target chromosomal region, of which six genes showed expression after stripe rust inoculation at the adult plant stage (Fig 3C). Integrated functional annotation revealed that four candidate genes potentially associated with plant disease resistance, such as *TraesCS2B01G162000* encoded an E3 ubiquitin ligase; *TraesCS2B01G163300* belonged to the multidrug and toxic compound extrusion (MATE) family; *TraesCS2B02G163600* was a member of the glycosyltransferase 8 family; and *TraesCS2B01G162100* encoded a Myb/SANT-like DNA-binding domain protein.

### 2.5. Interaction effects of resistance loci

The 153 F₅ RILs were classified into eight genotypic groups based on the molecular marker *csLV46G22, Xgwm533*, KASP_2BS for *Yr29*, *Yr30*, and *QYr.hzau-2BS*, respectively (Fig. 4; S3 Table). Analysis of the combined data showed that RILs carrying single resistance locus (*Yr29*, *Yr30*, and *QYr.hzau-2BS*) displayed significantly lower FDS compared to the lines lacking all resistance genes. *Yr29* was the most effective, conferring the lowest mean FDS of 38.20% (Fig. 4; S3 Table). The line carrying *QYr.hzau-2BS* had the mean FDS of 38.66%, whereas *Yr30* alone provided only a minor, but significant reduction (19.79%) on stripe rust. Among the two-gene combination, *Yr29* + *Yr30*, *Yr29*+ *QYr.hzau-2BS* and *Yr30* + *QYr.hzau-2BS* exhibited the mean FDS of 21.60%, 28.53% and 30.90% respectively. Lines carrying these three resistance genes showed significantly lower FDS than those carrying two. The combination *Yr29 + Yr30 + QYr.hzau-2BS* exhibited the lowest FDS (9.15%), approaching a near-immune state (Fig. 4; S3 Table).

**Fig 4.**
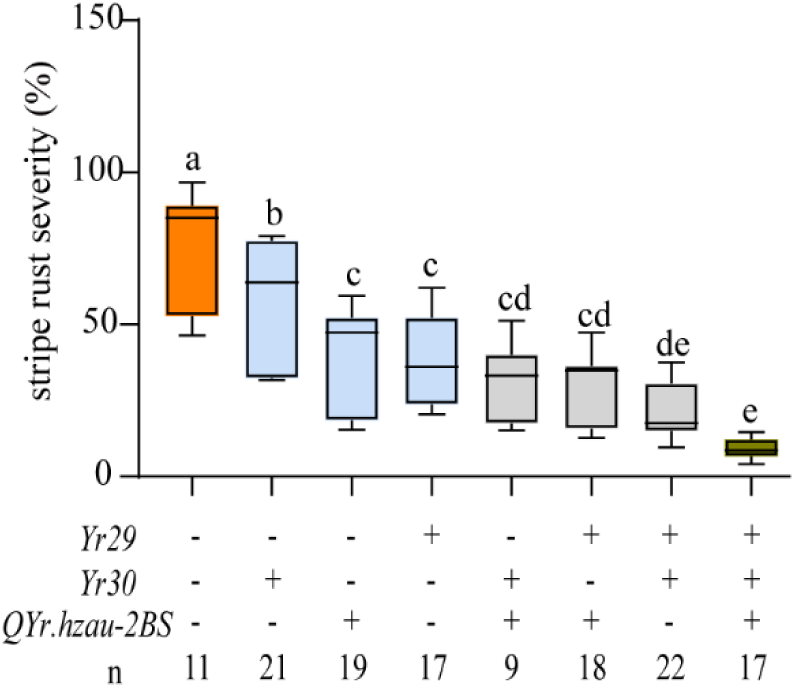
Effects of stacking resistance genes on stripe rust severity in ‘Apav#1’ × ‘Kijil’ RIL population. Box-and-whisker plots show the distribution of final disease severity among lines with different combinations of *Yr29*, *Yr30*, *QYr.hzau-2BS*. “−” indicates the absence of the QTL locus; whereas “+” indicates its presence. The center line, box, and whiskers showing the median, interquartile range and minimum to maximum values, respectively. One-way analysis of variance revealed significant differences (*F* [7, 48] = 7.121, *P* = 7.79 × 10^-6^). Groups annotated with different lowercase letters are significantly different at *P* < 0.05 (Duncan’s multiple range test). Different colors represent distinct genotype categories (no resistance genes, orange; one resistance gene, light blue; two resistance genes, gray; three resistance genes, yellow); “n” indicates the sample size of each combination.

## 3. Discussion

Developing wheat varieties with durable resistance remains a major objective of modern wheat breeding programs. In this study, Kijil, a CIMMYT-derived wheat line with superior agronomic performance, exhibited a high level of resistance to stripe rust across both Mexican and Chinese field environments. A total of five stable QTLs for APR to stripe rust were identified from Kijil, among these, the known resistance genes *Yr29* and *Yr30* were also confirmed based on the genotype and phenotype effect.

### 3.1. QYr.hzau-1BL (Yr29)

*QYr.hzau-1BL* was located in the interval between DArT markers 5410703 and 1862932, explaining 16.03–33.71% of the phenotypic variation. It was mapped to 671.27–676.07 Mb near the distal end of chromosome 1BL based on the Chinese Spring IWGSC RefSeq v1.0 (IWGSC 2018), where the well-known adult plant resistance gene *Lr46/Yr29/Sr58/Pm39/Ltn2* was closely linked to molecular marker *csLV46G22* (E. Lagudah, personal communication) and has been reported on this chromosome region within 2.6 Mb from *QYr.hzau-1BL* [18-20].*Yr29* has been deployed globally for over six decades and is known to provide partial APR to multiple diseases, including leaf rust and stripe rust. Its effect varies among genetic backgrounds, explaining 15.1–22.1% of the YR variation in ‘Pastor’ and 11.7–43.6% in ‘Francolin #1’ [20,21]. Similarly, *QYr.hzau-1BL* reduced YR severity by 16% and 21% in the ‘Borlaug 100’ [22] and Kijil backgrounds, respectively. The results confirmed that *Yr29* contributes stable, partial resistance across diverse environments and genetic backgrounds.

### 3.2. QYr.hzau-2BS

*QYr.hzau-2BS* was significantly associated with stripe rust resistance under Ciudad Obregón (Mexico) and Ezhou (China) rust environments, mapping between 134.55 Mb and 135.79 Mb on chromosome 2B (IWGSC RefSeq v1.0). Several stripe rust resistance genes have been mapped on 2BS chromosome, including *Yr5/Yr7/YrSp*, *Yr27*, *Yr31*, *Yr41*, *Yr43*, *Yr53*, and *Yr72*. *Yr27* originally derived from ‘Selkirk’ and it was closely linked with the leaf rust resistance gene *Lr13* [23], which is common in CIMMYT germplasm [24].

Recently, *Yr27* was cloned from a South African cultivar ‘Kariega’ by using a high-quality genome assembly method [25]. The gene spans 23.2 kb and encodes an NLR protein of 1,072 amino acids, sharing 97.3% sequence identity with *Lr13*. However, virulence to *Yr27* is now widespread, for example majority of Chinese races CYR32 and CYR33 as well as US races *PST*v-37 and *PST*v-52 were virulent to this gene and the virulence *Pst* race for *Yr27* was about 69.9% of 61 Chinese *Pst* isolates [26-28]. Despite this, *QYr.hzau-2BS* was consistently detected in Ezhou across multiple years in the present study, reducing mean disease severity by 12.61-18.13%. It was mapped 23.13 Mb away from *Yr27* and Kijil also lacked the gene-based molecular marker of *Yr27*. Thus, *QYr.hzau-2BS* should be a new stripe rust resistance locus from Kijil. In addition, we also found that it showed significant additive effect with *Yr29*, indicating its potential for gene pyramiding strategies to improve YR resistance in wheat breeding.

The KASP_2BS marker, developed for *QYr.hzau-2BS*, consistently differentiated between resistant and susceptible phenotypes across multiple environments. To elucidate the genetic basis of the quantitative trait locus *QYr.hzau-2BS* conferring resistance to stripe rust in wheat, we integrated public transcriptome data and screened genes within this interval based on expression patterns and functional annotation.

*TraesCS2B01G162000* encodes an E3 ubiquitin ligase, a class of enzymes that play pivotal regulatory roles in plant immunity by mediating the ubiquitination and degradation of key signaling components resulting in transducing defense responses [29,30]. Among these, the E3 ligase *TaVDIP1* mediates the ubiquitin-mediated degradation of the channel protein *TaVDAC1*, consequently inhibiting ROS accumulation and mitochondrial damage, which impairs the plant’s disease resistance [31]. Similarly, *TaGW2*, a RING-type E3 ligase, is induced by *Puccinia striiformis*. It suppresses ROS production by ubiquitinating and degrading the positive regulators *TaSnRK1γ* and *TaVPS24*, thereby enhancing wheat susceptibility [32]. In contrast, the C3HC4-type E3 ligase *RFEL1* stabilizes the negative regulator *TaNPR1* by ubiquitinating and degrading *TaNPR3*, activating the expression of downstream defense genes (such as *TaPR1* and *TaWRKY70*) and ultimately conferring broad-spectrum resistance to stripe rust, powdery mildew, and leaf rust in wheat [29]. This reveals the dual regulatory function of E3 ligases within the disease resistance network.

*TraesCS2B01G163300* belongs to the Multidrug and Toxic Compound Extrusion (MATE) transporter family. MATE proteins are primarily involved in the efflux of secondary metabolites, such as antimicrobial alkaloids or phenolics, and constitute an important component of chemical defense research has similarly demonstrated that MATE transporters constitute pivotal nodes within plant disease resistance networks. In *Arabidopsis thaliana*, for instance, they precisely regulate resistance by directly controlling the transport of core defense signaling molecules such as salicylic acid (such as *EDS5*, *ADS1*) [33]. Alternatively, they may impact on the expression of defense-related genes. In rice, overexpression of *OsMATE1/2* suppresses the expression of *PR1*, *PR5* and other genes, thereby exerting a potent negative regulatory effect on disease resistance [34].

*TraesCS2B01G163600* is a member of Glycosyltransferase Family 8 (GT8). Glycosyltransferase family 8 (GT8) comprises key enzymes involved in the biosynthesis and modification of plant cell wall polysaccharides, such as xylan [35]. Research indicates that expression levels of this family frequently fluctuate during pathogen infection, suggesting a potential role in disease resistance through enhancing cell wall structure or participating in defense signaling pathways. For example, *TaGT43H* and *TaIRX14*, belonging to the GT43 family in wheat, exhibit significantly upregulated expression following *Fusarium graminearum* infection. Their expression patterns correlate with plant resistance phenotypes, suggesting potential roles in cell wall remodeling or disease response [36]. However, direct causal relationships with disease resistance require further validation through loss-of-function or overexpression experiments. Overall, the GT8 family may play a crucial role in plant disease resistance mechanisms by regulating the synthesis of cell wall components such as xylan. Its specific functions and regulatory networks warrant further investigation to elucidate their mechanisms.

*TraesCS2B01G162100* encodes a protein containing a Myb/SANT-like DNA-binding domain. MYB (Myeloblastosis) transcription factors are one of the largest transcription factors families in plants and are widely involved in biotic and abiotic stress response, enhancing plant resistance by regulating the expression of downstream defense genes [37]. For example, in rice, *OsMYB30, OsMYB102* and *OsMYB108* directly activate defense-related genes to regulate resistance to rice blast and leaf blight [37,38]. In wheat, *TuMYB46L* forms a regulatory module with *TuACO3* to enhance resistance to powdery mildew (*Blumeria graminis* f. sp. *tritici*) by promoting ethylene synthesis [39]. These studies suggest that MYB transcription factors can play a central role in crop disease resistance through a multilevel regulatory network. However, the *TraesCS2B02G162100* of interest in this study differs from typical R2R3-MYB transcription factors (*OsMYB30*) in that its encoded protein contains a Myb/SANT-like structural domain (Pfam: pfam12776), indicating that this gene may belong to a functionally specialized subclass of the MYB superfamily, and its specific DNA-binding properties and regulatory functions need to be further investigated.

### 3.3. QYr.hzau-3BS (Yr30)

Chromosome 3BS is known as a hotspot for disease resistance genes, harboring *Fhb1* [40], *Yrns-B1* (*Yr30*) [8,41], *Sr2* [42], *Lr27* [43] and *Pm70* [44]. In this study, *QYr.hzau-3BS* mapped between DArT marker 1321522 and SSR marker *Xgwm533*, explaining 3.48–12.77% of phenotypic variation. Several APR stripe rust QTL on 3BS have been reported in varieties such as ‘Opata’ [8], ‘Kukri’ [45], and ‘Renan’ [46,47]. The closely linked molecular marker *Xgwm533* was also significantly associated with resistance in Apav#1/Kijil RIL population, suggesting that *QYr.hzau-3BS* corresponds to the *Yr30* region. Although their physical position overlapped, the additional validation—such as allelism tests or fine mapping—will be required to confirm their relationship.

### 3.4. QYr.hzau-3AS

*QYr.hzau-3AS* represents another APR locus detected exclusively in the Mexican test environments. Only a few YR resistance genes have been reported on chromosome 3AS, such as *YrTr1*, derived from the variety ‘Tres’, conferring ASR [48,49]. *Yr76* also provides ASR distinct from ‘Tyee’ and it was flanked by SSR markers *wmc11* and *wmc532* [50]. Lillemo et al. [51] identified an APR QTL on 3AS from ‘Saar’, and it was flanked by markers *Xstm844tcac* and *Xbarc310*, and this locus was only detected in Toluca environment. It was mapped to 7.14 Mb on the chromosome 3AS based on the Chinese Spring IWGSC RefSeq v1.0 (IWGSC 2018). Despite the 13 Mb difference in the physical location between *QYr.hzau-3AS* (19.32-20.23 Mb) and QTL mapped by Lillemo et al, the two loci may be the same due to both were detected only in Toluca, suggesting that this locus may confer race-specific or environment-specific resistance in the adult plant stage. However, the subsequent fine-tuning tools such as high-resolution genetic mapping, allelopathy tests, or haplotype analysis are still needed to confirm their relationship in future.

### 3.5. QYr.hzau-5DL

To date, no named YR resistance gene has been reported on chromosome 5DL, although several resistance QTL have been mapped. Suenaga et al. [47] identified an APR locus from ‘Oligoculm’ with flanking molecular marker of *Xwmc215* (472.37 Mb) on this chromosome, while Imtiaz et al. [52] and Liu et al. [53] mapped *QYr.nsw-5DL* and *QYr.caas-5DL* near 408.66 Mb. Another QTL, *QYr.GTM-5DL*, was reported in the Chinese landrace ‘Guangtoumai’ and it was flanked by *dCAPS-5722* at 449.29–451.17 Mb [54]. In contrast, *QYr.hzau-5DL*, identified in this study, was located at 337.69–361.65 Mb within the physical interval distinct from all previously reported loci. Although it explained a relatively small effect on stripe rust (3.75–10.97%), the QTL was consistently detected across Mexican and Chinese rust environment, indicating a stable and novel APR locus to YR.

### 3.6. Additive effects of resistance loci on stripe rust

Analysis of disease severities of RILs carrying combinations of resistance loci revealed that pyramiding multiple resistance genes is an effective strategy for enhancing resistance, which is consistent with previous reports [55]. The resistance gene *Yr29* not only significantly reduced stripe rust severity but also displayed a strong additive effect when combined with other resistance loci, leading to enhanced resistance. This conclusion is supported by several studies. For example, combining *Yr29* with *QYr.nwafu-4BL.3* reduced disease severity (DS) by 16.8–27.7% [56]. In the Chinese wheat cultivar Jimai 44, carrying the combination of *Yr29* and *YrJ44*, the mean DS was reduced to 29.5% [57]. Similarly, pyramiding *Yr29* with *Yr30* in variety Borlaug 100 reduced DS by 55% [22]. Although *Yr30* alone conferred a little detectable resistance in our population, it contributed to a pronounced synergistic effect in the three-gene combination with *Yr29* and *QYr.hzau-2BS*. The two-gene combination (*Yr29*+ *QYr.hzau-2BS*) achieved a DS of 28.53%, whereas the triple combination (*Yr29*+ *Yr30*+ *QYr.hzau-2BS*) further reduced DS to 9.15%, a level approaching immunity. This indicates that *Yr30*, despite its weak individual effect, is a key component within a polygenic resistance network, where it enhances the effects of other genes. These results collectively demonstrate that *Yr29* serves as a foundational component, effectively stacking with minor-effect genes such as *Yr30* and *QYr.hzau-2BS*. The generalizability of this stacking effect provides a strong theoretical basis for polygene pyramid and explains why *Yr29* is widely retained in global wheat breeding programs as a core trait for durable resistance.

In summary, we identified five stripe rust adult plant resistance (APR) loci in the Apav#1 × Kijil population. Two of these corresponded to the known genes *Yr29* and *Yr30*, whereas *QYr.hzau-2BS* and *QYr.hzau-5DL* are likely novel loci. A total of 4 potential candidate genes were predicted for the stable and major-effect resistance locus *QYr.hzau-2BS* and a KASP marker was developed for this locus to facilitate its utilization in wheat breeding. In addition, we also found the significant additive effects between detected resistance loci, especially for the gene combinations of *Yr29*. For example, the combination of *Yr29*, *Yr30*, and *QYr.hzau-2BS* formed a high-resistance core module. Therefore, it will be good if wheat breeders can use the related closely linked molecular markers to pyramid this core combination with other resistance loci for marker assisted selection, providing an effective strategy for developing wheat cultivars with enhanced and durable stripe rust resistance.

## 4. Materials and Methods

### 4.1. Plant materials

The CIMMYT wheat line ‘Apav#1’ (CIMMYT GID 1854090; pedigree: Avocet-*YrA*/Pavon) was susceptible to stripe rust at both seedling and adult plant stages, whereas Kijil (CIMMYT GID 6342979; pedigree: Klein Don Enrique*2/3/Fret2/Weebill1//Tacupeto F2001) was susceptible at the seedling stage but showed field resistance to the same *Pst* races at the adult plant stage. A population of 153 F₅ recombinant inbred lines (RILs) was developed from the cross between Apav#1 and Kijil at CIMMYT, using the method described by Yuan et al [58]. For field screening experiments, the Chinese wheat line ‘SY-You’ and a near-isoline ‘Avocet+*Yr31’* were used as susceptible controls and spreaders in China, while in Mexico, a mixture of Morocco and six susceptible lines derived from an ‘Avocet*’ × ‘*Attila*’* cross served as spreaders to ensure sufficient inoculum for phenotyping the RIL population.

### 4.2. Field trials and disease evaluation

The 153 RILs and their parents were evaluated for seedling resistance to YR in a greenhouse at Huazhong Agricultural University, China, using *Pst* races CYR33 and CYR34. Infection types (IT) were scored 12–14 days after inoculation based on a 0–9 scale [59]. The same materials, along with susceptible checks, were evaluated for adult plant resistance in multiple environments: Ciudad Obregón during 2015–2016 (YR2016Y), Toluca during 2016 and 2017 (YR2016T and YR2017T), El Batán in 2017 (YR2017B), and Ezhou City, Hubei Province, China, during 2019–2020, 2020–2021, and 2021–2022 (YR2020EZ, YR2021EZ, and YR2022EZ, respectively). Field experiments followed a randomized complete block design. In China, each line was sown in 1.5 m rows spaced 30 cm apart, with approximately 50 seeds per row, whereas in Mexico sowing was done as paired rows of 0.7 m length, 20 cm apart, on top of 80 cm wide raised beds. Forty RILs were randomly selected for two replications using an augmented design. A susceptible spreader row was planted alongside each plot.

In Toluca, Mexico, spreaders were inoculated at the jointing stage with *Pst* isolate Mex14.141, characterized by avirulence/virulence to *Yr1, 4, 5, 10, 15, 17, 24* and virulence to *Yr2, 3, 6, 7, 8, 9, 27, 31*, and *A* [60]. Natural infections incited epidemics at other two field sites in Mexico. In China, inoculations were conducted with a mixture of races CYR33 (avirulence/virulence formula based on seedling phenotypes: *Yr5, 8, 10, 15, 24, 26, 27, 32, Tr1/1, 6, 7, 9, 17, 18, 28, 29, 31, 43, 44, Exp2, SP*) [61] and CYR34 (*Yr1, 3, 4, H46, 5, 6, 15, 17, 18, 32, SP, Sd/2, 8, 9, 10, 12, 24[=26], 31, Su*) [62]. Inoculum was applied as a suspension of urediniospores in Soltrol 170 oil. YR disease severity (DS) was recorded using the modified Cobb’s Scale when susceptible checks reached ∼80% severity [63]. A second assessment was made one week later, when the checks reached 100% severity; these values were used as final disease severity (FDS) for analysis. FDS was recorded once at El Batán (YR2017B); twice at Ezhou during 2019-2021 (YR2020EZ, YR2021EZ) and once in 2022 (YR2022EZ); three times at Toluca during 2016-2017 (YR2016T) and once in 2015-2016 (YR2017T); and once at Ciudad Obregón (YR2016Y). Host response to infection was evaluated according to Roelfs et al following: R (resistant): small sized uredinia with necrosis [64]; MR (moderately resistant): small to intermediate sized uredinia with limited sporulation and visible chlorosis/necrosis; M (moderately resistant-moderately susceptible): medium sized uredinia with moderate sporulation and some chlorosis/necrosis; MS (moderately susceptible): medium sized uredinia with abundant sporulation and no chlorosis/necrosis; and S (susceptible): large uredinia with abundant sporulation and no chlorosis/necrosis.

Phenotypic data from multiple environments were combined using Best Linear Unbiased Prediction (BLUP), calculated as:

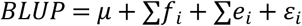

where μ is the population mean, *f₁* represents the genetic effect, *e₁* the environmental effect, and *ε₁* the random error [65].

### 4.3. Statistical analysis

The 153 RILs were classified into three categories following Singh and Rajaram [66]: homozygous parental-type susceptible (HPTS), homozygous parental-type resistant (HPTR), and OTHER (responses differing from both parents). The number of resistance genes was estimated using Mendelian segregation analysis [66,67]. A Chi-squared (χ²) test was used to assess goodness-of-fit to expected segregation ratios for two to five independent resistance loci. Correlation for FDS across environments were performed using IBM SPSS Statistics (Version 24) [68]. A one-way ANOVA for FDS across environments was performed using R software (version 4.5.2) [69], with statistical analyses tested with Levene’s test using the car package for variance homogeneity. Following a significant ANOVA, Duncan’s multiple range test (*agricolae* package) was used for post-hoc comparisons at α = 0.05 [70].

### 4.4. Genotyping, genetic mapping, and QTL analysis

Genomic DNA was extracted from the parents and approximately 20 seedlings per RIL using the CTAB method [71]. Genotyping was conducted using the DArT–GBS high-throughput platform (https://www.diversityarrays.com) at CIMMYT headquarter. A total of 84,300 presence/absence variation (PAV) markers and 51,348 SNP markers were obtained. After filtering for distorted segregation (*P* < 0.001), monomorphic loci, and markers with >20% missing data, 5,468 polymorphic markers were retained for linkage map construction. Genetic linkage maps were constructed using JoinMap 4.1 (Van Ooijen 2006) at a LOD threshold of 15.0 [72]. QTL analysis was performed using IciMapping 4.1 [73], based on FDS from each environment, and BLUP values obtained from multi-environmental data. For each locus, LOD scores, phenotypic variance explained (PVE), and additive effects were estimated. Significant QTLs (*P* < 0.05) were identified after 1,000 permutations, and MapChart (Voorrips 2002) was used to visualize linkage maps [74].

### 4.5. Molecular marker analysis

Both parents were genotyped with markers linked to 15 known *Yr* resistance genes (*Yr5/Yr7/YrSP, Yr9, Yr10, Yr15, Yr17, Yr18, Yr26, Yr27, Yr28, Yr29, Yr30*, *Yr36,* and *Yr46*. The primer sequences are in S2 Table. Among these, *csLV46G22* (E. Lagudah, personal communication) for *Yr29*, *KASPYr27* for *Yr27*, and *Xgwm533* for *Yr30* were polymorphic between two parents and subsequently used to genotype the entire RIL population [42].

Conventional PCR amplification protocols for the above genes were validated following Dreisigacker et al [71]. Each 10 μL reaction contained 1 μL genomic DNA (100 ng/μL), 5 μL 2× PCR Master Mix, 1 μL primer mix (forward and reverse), and 3 μL ddH₂O. The KASP PCR reactions (5 μL) included 1 μL DNA (100 ng/μL), 2.5 μL 2× KASP Master Mix, 0.1 μL FAM primer, 0.1 μL HEX primer, 0.2 μL common primer, and 1.1 μL water. The thermocycling profile was: pre-denaturation at 95°C for 5 min; 35 cycles of 95°C for 1 min, 58°C for 20 s, 72°C for 45 s; followed by final extension at 72°C for 10 min and hold at 4°C. PCR products amplified with *csLV46G22* were digested with *BspeI* endonuclease (37°C for 1 h) and resolved by 1.0% agarose gel electrophoresis. KASP markers were analyzed by real-time PCR (RT-PCR).

## Acknowledgments

We acknowledge Computation analysis performed on the bioinformatics computing platform at the National Key Laboratory of Crop Genetic Improvement, Huazhong Agricultural University. This work was supported by National Key Research and Development Program of China (grants 2024YFD1201105, 2022YFD1201300 and 2022YFD1201500), the Biological Breeding-National Science and Technology Major Project (2023ZD04025), National Natural Science Foundation of China (grant nos. W2412009, 32372173 and32501991), the Natural Science Foundation of Hubei Province of China (2024AFB154) and the Hubei Hongshan Laboratory (2022hspy001, 2021hskf008, and 2022hspy010). The work in Mexico was supported by the Australian Grains Research and Development Corporation (GRDC) with funding to the Australian Cereal Rust Control Program (ACRCP); and Accelerating Genetic Gains in Maize and Wheat (AGG) project Grant INV-003439 funded by the Bill and Melinda Gates Foundation (BMGF), the UK’s Foreign, Commonwealth and Development Office (FCDO).

## Supporting information

**S1 Fig.**
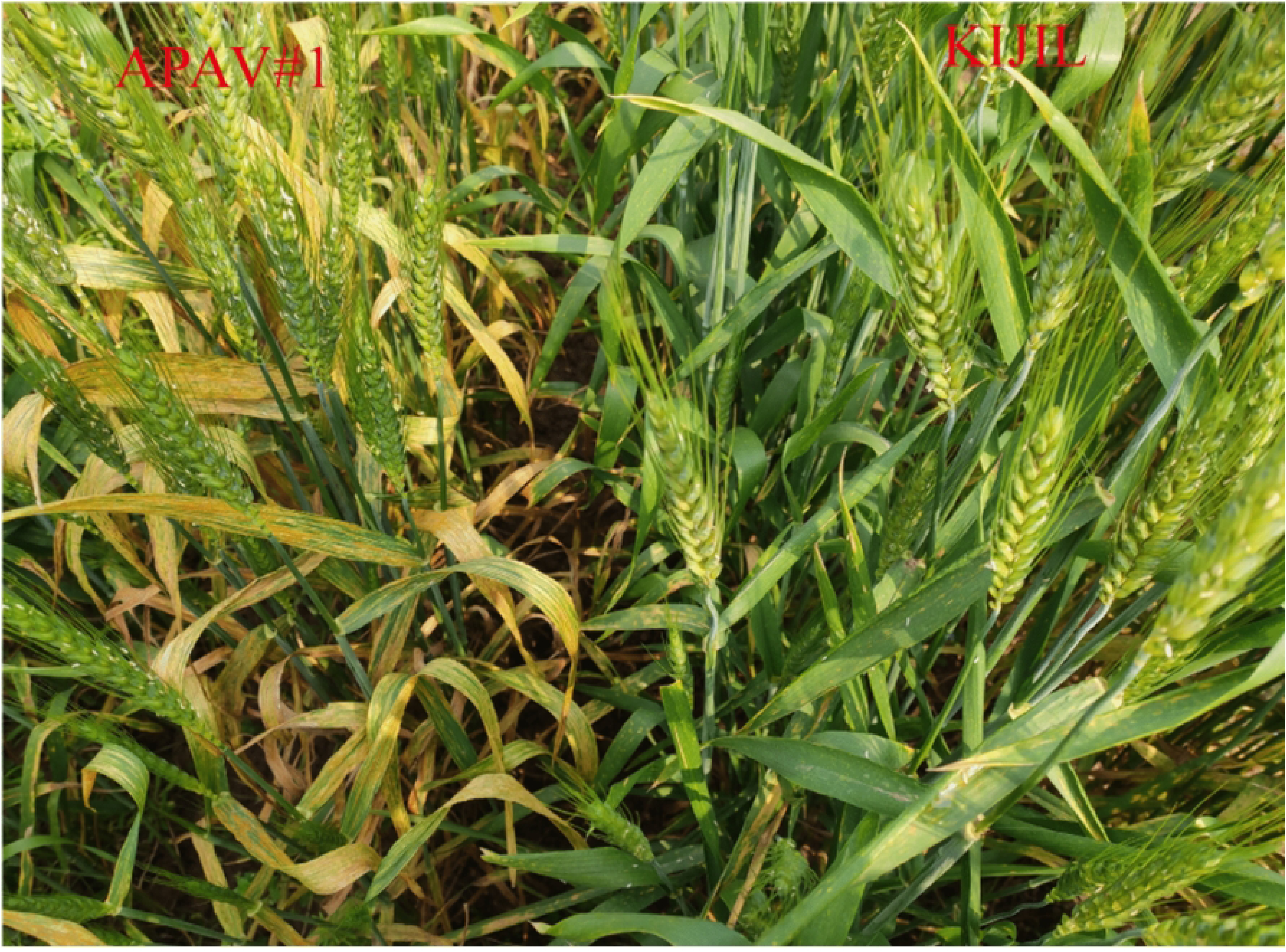
Field evaluation of stripe rust response. The susceptible parent ‘Apav#1’ (left) and the resistant parent ‘Kijil’ (right) are shown.

**S2 Fig.**
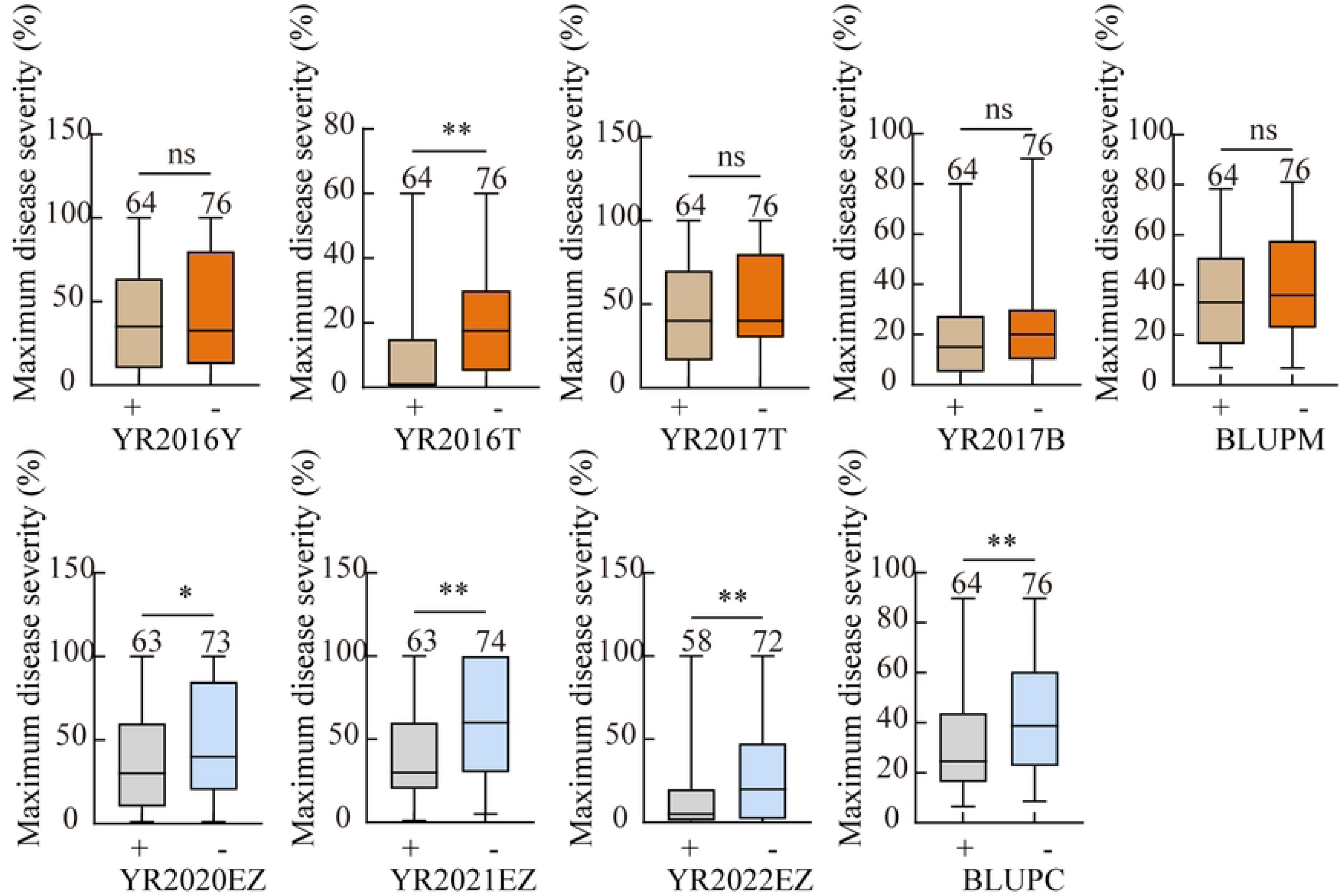
Association analysis for *QYr.hzau-2BS* using the KASP marker with stripe disease severity across nine environmrnts. The sample size was adequately designed. “−” indicates the absence of the QTL locus, whereas “+” indicates its presence. Significance levels are denoted by asterisks (*, *P* < 0.05; **, *P* < 0.01; ns = non-significant.).

**S1 Table. Phenotypic data (FDS) of the 153 RILs across nine environments used for QTL analysis**

**S2 Table. Primer sequences for the functional/closely-linked molecular markers of known resistance genes.**

**S3 Table. Phenotypic data (FDS) of different groups across seven environments used for analysis.**

**S1 File. Genotyping results of 5,468 molecular markers in the ‘Apav#1’× ‘Kijil’ RIL populatio**

